# Uncertainty and Exploration

**DOI:** 10.1101/265504

**Authors:** Samuel J. Gershman

**Author notes:** Address for correspondence: Samuel Gershman, Department of Psychology, Harvard University, 52 Oxford St., room 295.05, Cambridge, MA 02138.

## Abstract

In order to discover the most rewarding actions, agents must collect information about their environment, potentially foregoing reward. The optimal solution to this “explore-exploit” dilemma is often computationally challenging, but principled algorithmic approximations exist. These approximations utilize uncertainty about action values in different ways. Some *random* exploration algorithms scale the level of choice stochasticity with the level of uncertainty. Other *directed* exploration algorithms add a “bonus” to action values with high uncertainty. Random exploration algorithms are sensitive to *total* uncertainty across actions, whereas directed exploration algorithms are sensitive to *relative* uncertainty. This paper reports a multi-armed bandit experiment in which total and relative uncertainty were orthogonally manipulated. We found that humans employ both exploration strategies, and that these strategies are independently controlled by different uncertainty computations.

## Introduction

Uncertainty lies at the heart of decision making in the real world. A bee in search of nectar and a venture capitalist in search of an investment both need to explore their options in order to reduce their uncertainty, at the expense of exploiting the currently best option. In the absence of uncertainty, no explore-exploit dilemma would exist. The question addressed here is how humans use uncertainty to guide exploration.

The optimal solution to the explore-exploit dilemma is, except for some special cases (e.g., in foraging theory; Charnov, 1976; Stephens & Krebs, 1986), computationally intractable, but computer scientists have developed algorithmic approximations that can provably approach optimal behavior (Sutton & Barto, 1998). Psychologists have also studied algorithmic hypotheses about how humans balance exploration and exploitation (Cohen, McClure, & Yu, 2007; Hills et al., 2015; Mehlhorn et al., 2015), but only recently have the links between modern machine learning algorithms and psychological processes been systematically investigated (Gershman, 2018; Schulz, Konstantinidis, & Speekenbrink, 2017; Speekenbrink & Konstantinidis, 2015). Key to this synthesis is the idea that uncertainty can guide exploration in two qualitatively different ways: by adding *randomness* into choice behavior, or by directing choice towards uncertain options.

The pioneering work of Wilson, Geana, White, Ludvig, and Cohen (2014) demonstrated that humans use both random and directed exploration in a carefully designed two-armed bandit task. When the subject had more experience with one option (and hence less uncertainty), she favored the more uncertain option, indicating a form of directed exploration. In addition, subjects increased the randomness in their choices when they were more uncertain, spreading their choices across both high- and low-value options. Directed and random exploration strategies are dissociable, developing on different timescales across the lifespan (Somerville et al., 2017) and relying on different neural substrates (Zajkowski, Kossut, & Wilson, 2017).

The directed/random distinction is closely related to the distinction between two families of exploration algorithms that have been studied extensively in machine learning. Directed exploration can be realized by adding *uncertainty bonuses* to estimated values (Auer, Cesa-Bianchi, & Fischer, 2002; Brafman & Tennenholtz, 2002; Dayan & Sejnowski, 1996; Kolter & Ng, 2009; Srinivas, Krause, Seeger, & Kakade, 2010). One of the most prominent versions of this approach is known as the *upper confidence bound* (UCB) algorithm (Auer et al., 2002), which chooses action *a*_*t*_ on trial *t* according to:

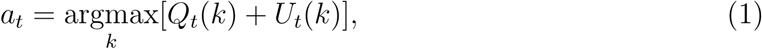

where *k* indexes actions and *U*_*t*_(*k*) is the upper confidence bound that plays the role of an uncertainty bonus. Under a Bayesian analysis (Srinivas et al., 2010), *Q*_*t*_(*k*) corresponds to the posterior mean and the uncertainty bonus is proportional to the posterior standard deviation *σ*_*t*_(*k*).

Random exploration has an even older pedigree in machine learning, dating back to Thompson’s work in the 1930s (Thompson, 1933). In what is now known as *Thompson sampling*, a random value function from the posterior is drawn and then the agent chooses greedily with respect to the random draw. Like UCB, Thompson sampling uses uncertainty to promote exploration, but does so by encouraging stochasticity (i.e., a form of probability matching) rather than via a response bias.^1^

In psychophysical terms, uncertainty in Thompson sampling changes the slope of the function relating action values to choice probabilities, whereas uncertainty in UCB changes the intercept (indifference point).^2^ The role of uncertainty in Thompson sampling can be understood by recognizing that sampling from a distribution over values implies that variability in this distribution directly translates to variability in choices. By contrast, the role of uncertainty in UCB can be understood by recognizing that the uncertainty bonus acts in the same way as a boost in reward, shifting choice probabilities towards the more uncertain option without altering the level of choice stochasticity.

Recognizing the dissociable (and possibly complementary) nature of directed and random exploration, computer scientists have also constructed hybrids of UCB and Thompson sampling (Chapelle & Li, 2011; May, Korda, Lee, & Leslie, 2012). A recent report (Gershman, 2018) provided the most direct evidence for such hybrids in human decision making, demonstrating that uncertainty influences both the intercept and slope of the choice probability function. Critically, the intercept and slope effects derive from different uncertainty computations: the intercept is governed by *relative* uncertainty (the difference in posterior uncertainty between the two options, defined formally below), whereas the slope is governed by *total* uncertainty (the sum of posterior uncertainty across the options). This suggests that experimental manipulations of these two factors should produce dissociable effects.

Here we pursue this line of reasoning using a two-armed bandit task, with the additional twist that we inform subjects about the riskiness of each arm (Figure 1). On each trial, participants were given a choice between two arms, labeled either as “safe” (S) or “risky” (R). The safe arms always delivered the same reward, whereas the risky arms delivered stochastic rewards (with Gaussian noise). Denoting trial types by compound labels (e.g., “SR” denote trials in which the left arm is safe and the right arm is risky), we used the comparison between preference for arm 1 on SR and RS trials to isolate the effects of relative uncertainty, holding total uncertainty fixed, and the comparison between preference for arm 1 on SS and RR trials to isolate the effects of total uncertainty, holding relative uncertainty fixed (Figure 2). By independently manipulating relative uncertainty (SR vs. RS) and total uncertainty (SS vs. RR), we can go beyond the correlational findings of Gershman (2018) to causally test the predictions of UCB and Thompson sampling. In particular, we predicted that the SS condition would increase the slope of the choice probability function (affecting random but not directed exploration) relative to the RR condition, whereas the RS condition would shift the intercept of the choice probability function (affecting directed but not random exploration) relative to the SR condition.

**Figure 1.**
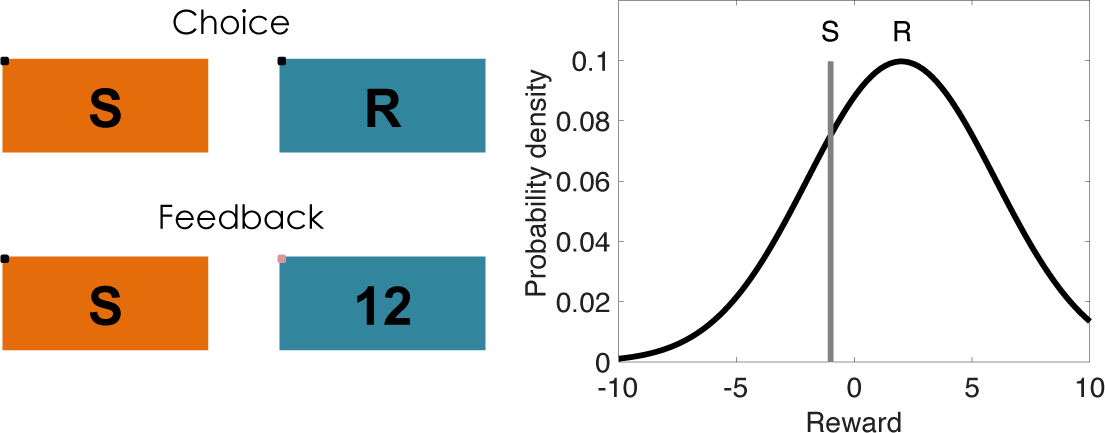
Task design. (Left) On each trial, subjects choose between two options and receive reward feedback in the form of points. Safe options are denoted by “S” and risky options are denoted by “R.” On each block, one or both of the options may be safe or risky. (Right) The rewards for risky options are drawn from a Gaussian distribution that remains constant during each block. The rewards for safe options are deterministic. Both the mean for the risky option and the reward value of the safe option are drawn from a zero-mean Gaussian distribution that is resampled at each block transition.

**Figure 2.**
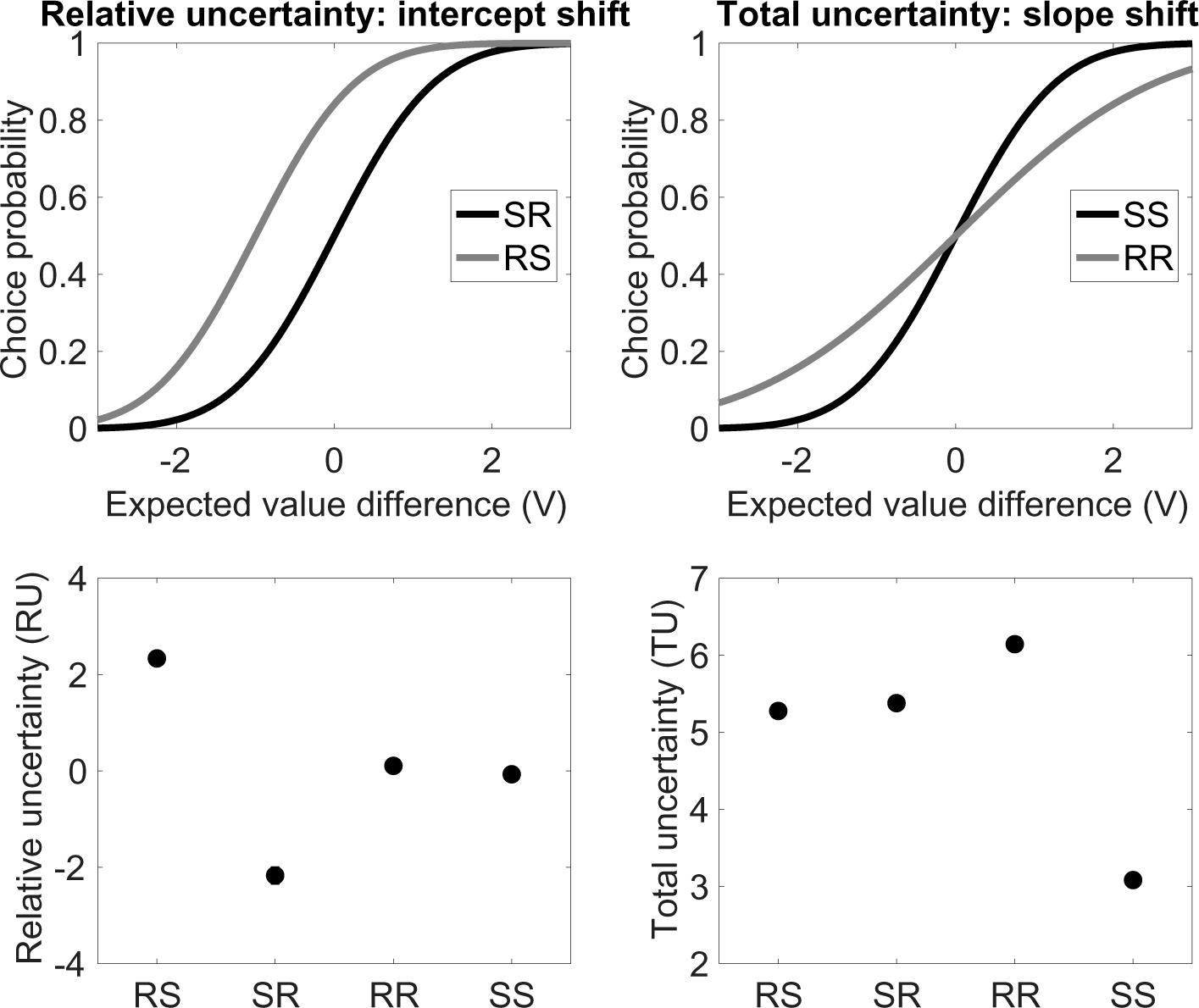
Relative and total uncertainty. (Top) Illustration of how the probability of choosing option 1 changes as a function of the experimental condition and form of uncertainty. V represents the difference between the expected value of option 1 and the expected value of option 2. (Bottom) Average relative and total uncertainty in each condition. Conditions are denoted by safe/risky compounds; for example, “SR” denotes a trial in which option 1 is safe and option 2 is risky.

## Materials and Methods

Code and data for reproducing all analyses reported in this paper, as well as Javascript code for re-running the experiments, are available at https://github.com/sjgershm/exploration_uncertainty.

## Subjects

Forty-six subjects were recruited via the Amazon Mechanical Turk web service and paid $2.00. The sample size was chosen to be comparable to previous studies using a similar experimental paradigm (Gershman, 2018). The experiments were approved by the Harvard Institutional Review Board.

### Stimuli and Procedure

Participants played 30 two-armed bandits, each for one block of 10 trials. On each trial, participants chose one of the arms and received reward feedback (points). They were instructed to choose the “slot machine” (corresponding to an arm) that maximizes their total points. On each block, the mean reward *μ*(*k*) for each arm was drawn from a Gaussian with mean 0 and variance 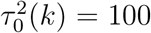. The arms were randomly designated “safe” or “risky,” indicated by an S or R (respectively), and these designations were randomly resampled after a block transition. When participants chose the risky arm, they received stochastic rewards drawn from a Gaussian with mean *μ*(*R*) and variance *τ*^2^(*R*) = 16. When participants chose the safe arm, they received a reward of *μ*(*S*).

The exact instructions for participants were as follows:

> In this task, you have a choice between two slot machines, represented by colored buttons. When you click one of the buttons, you will win or lose points. Choosing the same slot machine will not always give you the same points, but one slot machine is always better than the other. Your goal is to choose the slot machine that will give you the most points. After making your choice, you will receive feedback about the outcome. Sometimes the machines are “safe” (always delivering the same feedback), and sometimes the machines are “risky” (delivering variable feedback). Before you make a choice, you will get information about each machine: “S” indicates SAFE, “R” indicates RISKY. Note that safe/risky is independent of how rewarding a machine is: a risky machine may deliver more points on average than a safe machine, and vice versa. You cannot predict how good a machine is based on whether it is safe or risky. You will play 30 games, each with a different pair of slot machines. Each game will consist of 10 trials.

### Belief updating model

To derive estimates of expected value and uncertainty, we assume that subjects approximate an ideal Bayesian learner. Given the Gaussian distributional structure underlying our task, the posterior over the value of arm *k* is Gaussian with mean *Q*_*t*_(*k*) and variance 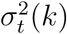. These sufficient statistics can be recursively updated using the Kalman filtering equations:

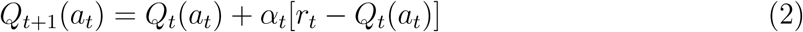

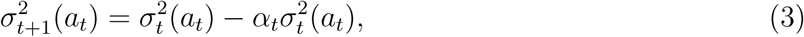

where *a*_*t*_ is the chosen arm, *r*_*t*_ is the received reward, and the learning rate *α*_*t*_ is given by:

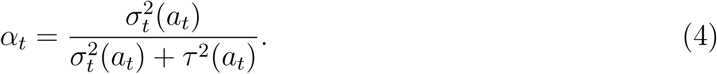

Note that only the chosen option’s mean and variance are updated after each trial. The initial values were set to the prior means, *Q*_1_(*k*) = 0 for all *k*, and the initial variances were set to the prior variance, 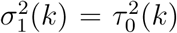. The value of *τ*^2^ was set to 16 (its true value) for risky options, and to 0.00001 for safe options (to avoid numerical issues, the variance was not set exactly to 0). Although the Kalman filter is an idealization of human learning, it has been shown to account well for human behavior in bandit tasks (Daw, O’doherty, Dayan, Seymour, & Dolan, 2006; Gershman, 2018; Schulz, Konstantinidis, & Speekenbrink, 2015; Speekenbrink & Konstantinidis, 2015).

### Choice probability analysis

Gershman (2018) showed that Thompson sampling, UCB, and a particular hybrid of the two imply a probit regression model of choice:

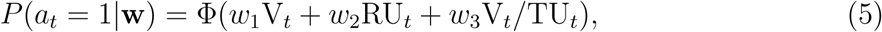

where ϕ(·) is the cumulative distribution function of the standard Gaussian distribution (mean 0 and variance 1), and the regressors are defined as follows:

- Estimated value difference, V_*t*_ = *Q*_*t*_(1) - *Q*_*t*_(2).
- Relative uncertainty, RU_*t*_ = *σ*_*t*_(1) - *σ*_*t*_(2).
- Total uncertainty, 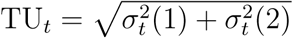.

As shown in Gershman (2018), Thompson sampling predicts a significant positive effect of V/TU on choice probability, but not of RU or V, whereas UCB predicts a significant positive effect of both V and RU, but not of V/TU (Figure 2, top left). We used maximum likelihood estimation to fit the coefficients (w) in the probit regression model.

In addition to this model-based analysis, we analyzed choices as a function of experimental condition (i.e., RS, SR, RR, SS).

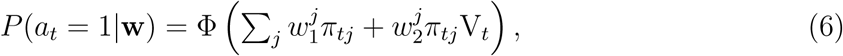

where π_*tj*_ = 1 if trial *t* is assigned to condition *j*, and 0 otherwise. We refer to the *w*_1_ terms as intercepts and the *w*_2_ terms as slopes.^3^

### Response time analysis

We examined response times as an additional source of evidence about exploration strategies. Our hypotheses are motivated by a sequential sampling framework, according to which the value difference between two options drives a stochastic accumulator until it reaches a decision threshold (Busemeyer & Townsend, 1993; Krajbich, Armel, & Rangel, 2010; Milosavljevic, Malmaud, Huth, Koch, & Rangel, 2010; Ratcliff & Frank, 2012; Summerfield & Tsetsos, 2012). Formally, the decision variable μ_*t*_(*τ*) evolves over time (within a trial, index by *τ*) as follows:

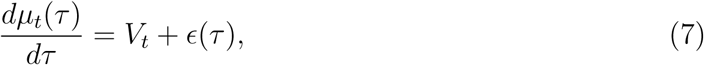

where *V*_*t*_ (the value difference between options 1 and 2 defined above) is a deterministic “drift” term, and ϵ(*τ*) is a stochastic “diffusion” term. Without loss of generality, it is conventional to assume that ϵ(*τ*) is drawn from a standard Gaussian distribution (mean of 0 and variance of 1). The decision variable evolves until it hits one of two thresholds ±*B* (corresponding to the two options), at which point a decision is made. If we assume, following the logic of the previous section, that both UCB and Thompson sampling are implemented by a linear transformation of the value difference (see Gershman, 2018, for more details), then the decision variable will evolve according to:

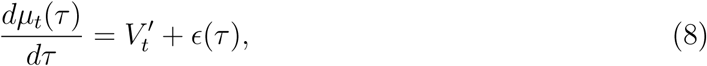

where 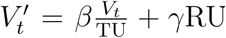, and *β* and *γ* are coefficients controlling the relative contribution of random and directed exploration, respectively. Under this model, the expected response time is given analytically by:

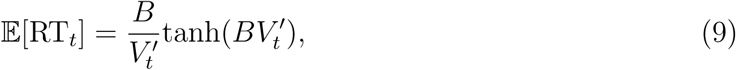

where tanh is the hyperbolic tangent function. Since the expected response time is a monotonically decreasing function of 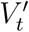, it decreases with RU and increases with TU (see also Busemeyer & Townsend, 1993).

## Results

The hybrid random/directed exploration model hypothesizes that all three computational regressors (V, RU, V/TU) should be predictors of choice. We therefore confirmed that maximum likelihood estimates of the corresponding coefficients were significantly greater than 0 [Figure 3; V: *t*(13797) = 16.48, *p* < 0.0001; RU: *t*(13797) = 4.9, *p* < 0.0001; V/TU: *t*(13797) = 6.04, *p* < 0.0001]. Model comparison using the Bayesian and Akaike information criteria strongly favored this three-parameter model over a one-parameter model with only the V regressor, a one-parameter model with V/TU, and a two-parameter model with V and RU (Table 1). This supports previous results showing that humans use both random and directed exploration strategies (Gershman, 2018; Somerville et al., 2017; Wilson et al., 2014).

**Figure 3.**
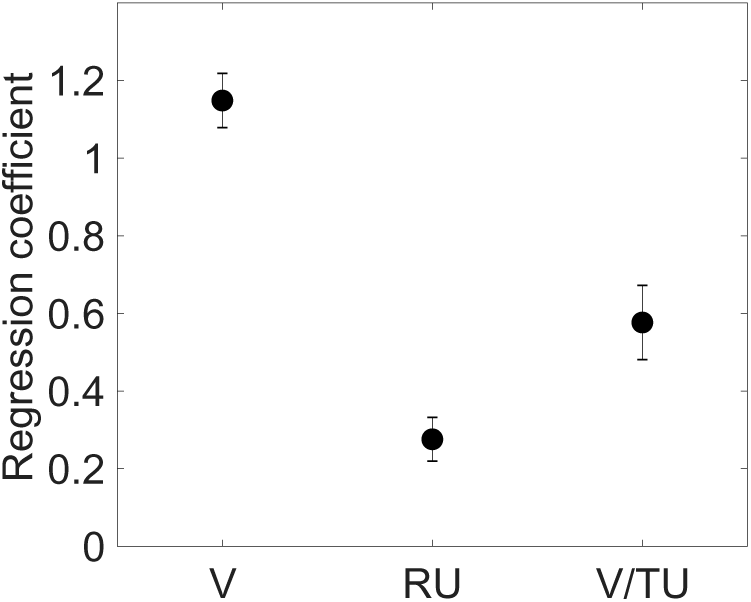
Probit regression results: computational variables. Fixed effects parameter estimates and standard errors for each regressor. V = estimated value difference between irms, RU = relative uncertainty, TU = total uncertainty (see Materials and Methods for etails).

**Table 1:** Model comparison results. BIC = Bayesian information criterion; AIC = Akaike information criterion. Lower values indicate higher model evidence.

| Model | BIC | AIC |
| --- | --- | --- |
| V | 13493 | 13478 |
| V/TU | 17262 | 17247 |
| V + RU | 12874 | 12836 |
| V + RU + V/TU | 12689 | 12621 |

We next addressed the central question of the paper: can random and directed exploration be independently manipulated? As predicted, changing relative uncertainty (RS vs. SR) alter the intercept of the choice probability function (Figure 3): there was a significant difference between the intercepts for the RS and SR conditions [*F*(1,13792) = 13.92, *p* < 0.001]. Furthermore, the RS intercept was significantly greater than 0 [*t*(13792) = 2.33, *p* < 0.02] and the SR intercept was significantly less than 0 [*t*(13792) = 4.47, *p* < 0.001], indicating an uncertainty-directed choice bias, as predicted by the theory. In other words, uncertainty boosted the value of an arm, shifting choice probability towards that arm. Critically, there was no significant effect of total uncertainty [RR vs. SS, *p* = 0.40], consistent with the hypothesis that uncertainty-directed biases only emerge when there is a difference in relative uncertainty.

The pattern is flipped when we inspect the slope parameter estimates (Figure 4): increasing total uncertainty (RR vs. SS) reduced the slope [*F*(1,13792) = 20.12, *p* < 0.0001], but relative uncertainty (RS vs. SR) did not have a significant effect (*p* = 0.72). This finding is consistent with our hypothesis that the random component of exploration would be specifically sensitive to changes in total uncertainty. In other words, the decreased slope for RR indicates greater choice stochasticity when total uncertainty was higher. An alternative possibility is that the increased level of payoff variance in RR compared to SS causes value differences to be more strongly driven by payoff noise. In other words, even if subjects were not using random exploration, they might still show a difference in choice behavior between the RR and SS conditions. However, this hypothesis cannot account completely for the RR vs. SS effect, because our analysis quantifies how choice probability differs between conditions for the same estimated value difference. Thus, even if the conditions differed in their effects on value learning, this analysis controls for such differences.

**Figure 4.**
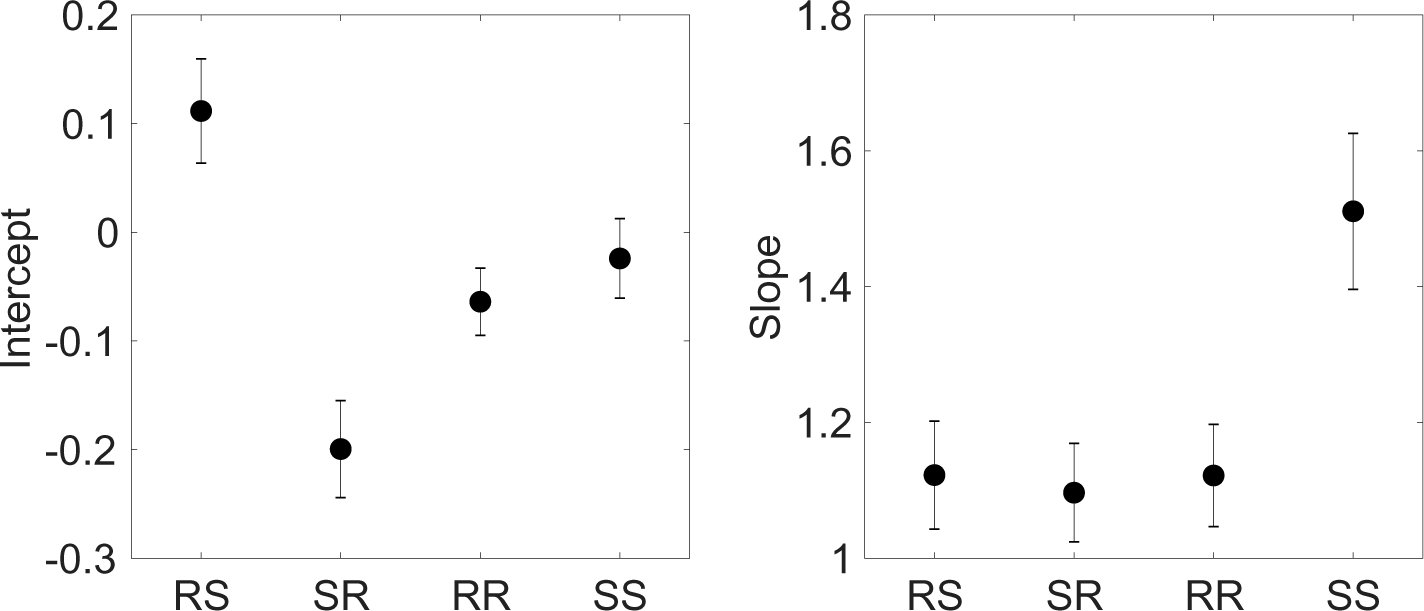
Probit regression results: experimental variables. Fixed effects parameter estimates and standard errors for each regressor. (Left) Intercept coefficients. (Right) Slope coefficients.

Sequential sampling models have also been shown to jointly predict choice and response time in reinforcement learning tasks, where the value is dynamically updated based on experience (Frank et al., 2015; Millner, Gershman, Nock, & den Ouden, 2018; Pedersen, Frank, & Biele, 2017). Within the sequential sampling framework, a directed exploration strategy predicts that response times should be faster for risky choices than for safe choices on RS and SR trials, because uncertainty acts as a bonus added to the values (see Materials and Methods). This prediction was confirmed in our data [Figure 5; *t*(45) = 2.01, *p* < 0.05].

**Figure 5.**
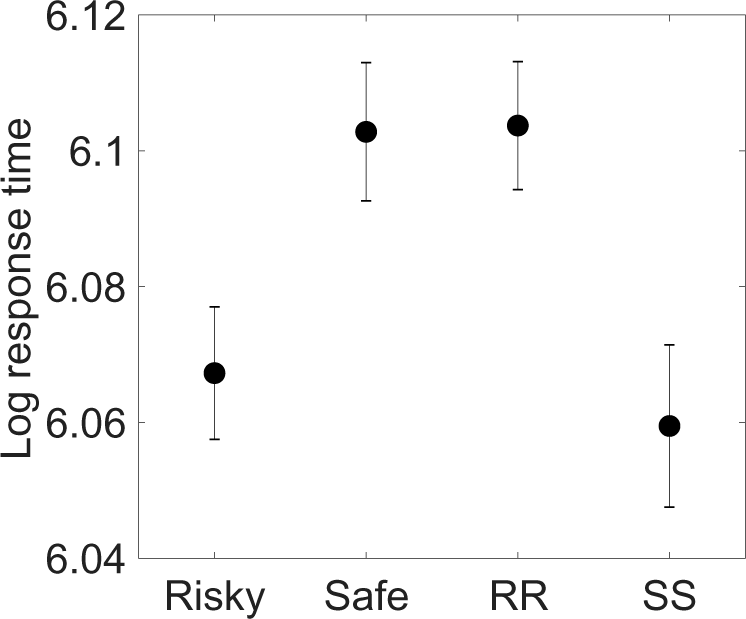
Response time analysis. Log response times (mean ± within-subject standard error). “Risky” denotes risky choices on SR and RS trials; “Safe” denotes safe choices on SR and RS trials. SS and RR response times are collapsed across arms.

When uncertainty is equated across the arms, directed exploration does not predict any difference in response time as a function of total uncertainty. Random exploration strategies, in contrast, correctly predict that total uncertainty will act divisively on values, thereby slowing response times when both arms are risky compared to when both arms are safe [Figure 5; *t*(45) = 2.28, *p* < 0.05].

## Discussion

By separately manipulating relative and total uncertainty, we were able to independently influence directed and random exploration, lending support to the contention that these strategies coexist and jointly determine exploratory behavior (Gershman, 2018; Wilson et al., 2014). In particular, increasing relative uncertainty by making one option riskier than the other caused participants to shift their preference towards the risky option in a value-independent manner, consistent with a change in the intercept (indifference point) of the choice probability function. This manipulation had no effect on the slope of the choice probability function. In contrast, increasing total uncertainty by making both options risky decreased the slope relative to when both options were safe, with no effect on the intercept. In other words, increasing total uncertainty caused choices to become more stochastic.

This dissociation between strategies guiding choice behavior was mirrored by a dissociation in response times. When one option was riskier than the other, subjects were faster in choosing the risky option. However, the same subjects were *slower* when both options were risky compared to when both options were safe. Taken together, these findings demonstrate that risk can have qualitatively different effects on both choice and response time, depending on the underlying uncertainty computation (i.e., total vs. relative). Consistent with our findings, previous work has shown that payoff variance increases exploration (Lejarraga, Hertwig, & Gonzalez, 2012; Wulff, Mergenthaler-Canseco, & Hertwig, 2018). However, this work did not distinguish between random and directed exploration.

We note here that preference for the risky option appears to be in opposition to what would be predicted by theories of risk aversion, as pointed out by previous authors (Gershman, 2018; Payzan-LeNestour & Bossaerts, 2012; Wilson et al., 2014). This does not mean that our subjects were not risk averse; rather, we suspect that the imperative to explore in a multi-armed bandit task overwhelmed the tendency to avoid risks seen in typical gambling paradigms.

The role of uncertainty-guided exploration has come to occupy an increasingly important place in theories of reinforcement learning (Gershman & Niv, 2015; Knox, Otto, Stone, & Love, 2011; Navarro, Newell, & Schulze, 2016; Payzan-LeNestour & Bossaerts, 2011; Pearson, Hayden, Raghavachari, & Platt, 2009; Schulz et al., 2015; Speekenbrink & Konstantinidis, 2015; Zhang & Yu, 2013), superseding earlier models of exploratory choice based on a fixed source of decision noise, as in 𝜖-greedy and softmax policies (e.g., Daw et al., 2006). This shift has been accompanied by a deeper understanding of how reinforcement learning circuits in the basal ganglia compute, represent, and transmit uncertainty to downstream decision making circuits (Gershman, 2017; Lak, Nomoto, Keramati, Sakagami, & Kepecs, 2017; Starkweather, Babayan, Uchida, & Gershman, 2017). Despite this progress, we are only beginning to understand how these circuits implement dissociable channels regulating directed and random exploration (Warren et al., 2017; Zajkowski et al., 2017). Furthermore, a number of inconsistencies exist in the literature; for example, some studies do not find evidence for uncertainty bonuses in exploration (Daw et al., 2006; Riefer, Prior, Blair, Pavey, & Love, 2017). Resolving these inconsistencies will require a more complete picture of the different factors governing directed and random exploration, including individual differences, internal states (e.g., stress, cognitive load), and task structure.

## Acknowledgments

I am grateful to Eric Schulz for helpful discussions. This research was supported by the NSF Collaborative Research in Computational Neuroscience (CRCNS) Program Grant R01-1207833 and the Office of Naval Research under grant N000141712984.

## Footnotes

1 The idea that uncertainty promotes choice stochasticity is present in some theories of decision making, notably Decision Field Theory (Busemeyer & Townsend, 1993). Thompson sampling differs formally in that stochasticity is driven by posterior uncertainty (see the formal description below), whereas in Decision Field Theory it is driven by payoff variance. Typically uncertainty and payoff variance are correlated, a fact that we exploit in our experimental design (see also Leuker, Pachur, Hertwig, & Pleskac, 2018).

2 We have assumed here, following Gershman (2018), that some noise is added to the values that enter into the UCB computation. Without this assumption (or some other sources of stochasticity), we would not be able to capture variability in choices.

3 Note that although this analysis is “model-free” in the sense that it does not use the computationally derived uncertainty regressors, it is still dependent on the model-based value estimates.

